# Linker dependence of avidity in multivalent interactions between disordered proteins

**DOI:** 10.1101/625327

**Authors:** Charlotte S. Sørensen, Agnieszka Jendroszek, Magnus Kjaergaard

## Abstract

Multidomain proteins often interact through several independent binding sites connected by disordered linkers. The architecture of such linkers affect avidity by modulating the effective concentration of intra-molecular binding. The linker dependence of avidity has been estimated theoretically using simple physical models, but such models have not been tested experimentally since the effective concentrations could not be measured directly. We have developed a model system for bivalent protein interactions connected by disordered linkers, where the effective concentration can be measured using a competition experiment. We characterized the bivalent protein interactions kinetically and thermodynamically for a variety of linker lengths and interaction strengths. In total, this allowed us to critically assess the existing theoretical models of avidity in disordered, multivalent interactions. As expected, the onset of avidity occurs when the effective concentration reached the dissociation constant of the weakest interaction. Avidity decreased monotonously with linker length, but only by a third of what is predicted by theoretical models. We suggest that the length dependence of avidity is attenuated by compensating mechanisms such as linker interactions or entanglement. The direct role of linkers in avidity suggest they provide a generic mechanism for allosteric regulation of disordered, multivalent proteins.

## Introduction

Protein-protein interactions often consist of several independent interactions that form simultaneously. Compared to monovalent interactions, such multivalent interactions have several functional advantages: Monovalent interactions tend to be all-or-nothing, while multivalent interactions can have different binding modes with different affinities.^1^ Multivalency also allows weak interactions to collectively form much stronger interactions, a phenomenon known as avidity.^2,3^ Multivalency and avidity are particularly important in signalling networks, where weak and modular protein-protein interactions organize signaling molecules in space.^4,5^

Multivalent interactions often occur through intrinsically disordered regions that contain short protein interaction motifs. These motifs are either known as SLiMs, short linear motifs,^6^ or MoRF, molecular recognition features.^7^ Multivalency often requires the segments surrounding the interaction site to be disordered to allow the protein to contact several binding sites.^8^ The affinity of such sites is often limited due to their small size, and hence multiple MoRFs are often combined into a multivalent interaction that are strengthened by avidity.^1,9^ Avidity is thus crucial in the interactions of disordered proteins, but little work has addressed the role of the connecting linker in determining avidity. For multivalent MoRFs this connection can typically be expressed in terms of the length and physical characteristics of the sequence separating the binding sites. The linker region is thus likely to determine how much an additional interaction adds to the total affinity, and at which point an interaction become so weak that it no longer contributes avidity enhancement.

Avidity has been investigated thoroughly both theoretically and experimentally. Jencks proposed that the free energy of binding in a bivalent interaction could be decomposed into contributions from each of the individual interactions and a connection free energy, which is mainly entropic (Fig. 1A).^10^ The connection energy is determined by the connection between the molecules and adds a constant free energy regardless of the strength of the individual interactions. Subsequently, much effort has been spent on estimating the connection energy from the topology of the interaction sites.^11–13^ Alternatively, avidity can be described in kinetic terms as sketched in Fig. 1A for a bivalent interaction. Following the initial binding of one site, the second binding step occurs intra-molecularly. The rate of this reaction depends both on the intermolecular rate constant and the effective concentration. The effective concentration, c_eff_, contains the contributions from the linker. An advantage of formulating avidity in terms of effective concentrations is that it can be measured independently of the overall binding reaction through competition experiments.^14,15^ In disordered proteins, effective concentration are practically never measured experimentally, but are typically estimated from polymer models. For a bivalent interaction, the avidity binding constant has often successfully been expressed as:^16,17^

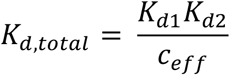

**Figure 1:**
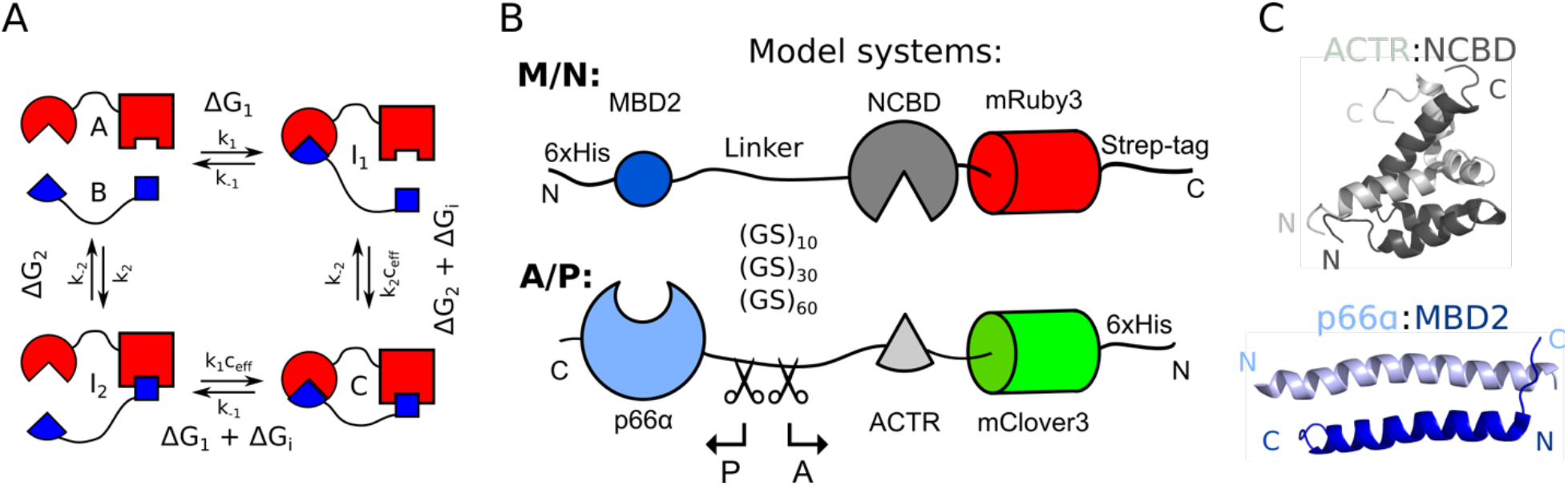
Design of a model system for bivalent interactions. Schematic illustration of bivalent interaction between A and B with two non-identical binding sites is shown in (A). Association of A and B can lead to formation of two types of monovalent complexes: I_1_ with association and dissociation rates k_1_ and k_-1_ respectively and free energy of interaction ΔG_1_ or I_2_ with k_on_ and k_off_ rates k_2_ and k_-2_ respectively and free energy of interaction ΔG_2_. Initial intermolecular reaction can be followed by binding of the second binding site and formation of bivalent complex C. Rate of this intramolecular reaction depends on the rate of intermolecular reaction (k_1_ or k_2_) and effective concentration (c_eff_), whereas free energy is expressed as a sum of the free energy of individual interaction (ΔG_1_ or ΔG_2_) and connection Gibbs energy ΔG_i_. (B) Model system used in the study contains two interacting proteins: M/N and A/P, each with two non-identical binding sites (MBD2 and NCBD or p66α and ACTR) separated by the linkers of 10, 30 or 60 GS repeats, a fluorescent domain located in the proximity of the weaker interacting pair (NCBD and ACTR) and tags: 6×His-tag used for purification and SA-Strep-tag used for both purification and oriented capture of M/N on the SPR chip. Monovalent interactions are studied using constructs of A/P, where each protein have 15 residues of the GS linker. (C) NMR structures of the interacting pairs used in the study: ACTR:NCBD (PDB: 1KBH) and MBD2:p66α (PDB: 2L2L).

This is equivalent to adding a constant coupling term to the free energies of the individual interactions. This model, however, is an approximation that will break down when two populations of the singly bound species (I_1_ and I_2_ in Fig 1A) cannot be ignored. In practice, this assumption has worked well as most studies of avidity has been motivated by the development of multivalent drugs: A high affinity inhibitor can be created by stringing together inhibitors of lower affinity, and studies of the connection has typically aimed at maximizing avidity.^2,18,19^ Therefore, previous studies used short linkers and strong binding partners. These conditions do not apply to interactions between multivalent, disordered proteins, where the linking architecture may be long and complex and individual interactions may be weak. To understand the role of avidity in multivalent protein interactions, we thus need to study it under conditions typical of those encountered in molecular biology.

The role of linking architecture in avidity may best be explored using a model system mimicking conditions relevant for multivalent MoRFs. We thus developed a model system for bivalent interactions between disordered proteins, where the connecting linker and the interaction strengths could be varied and measured independently. This model system allowed us to confirm that the onset of avidity occurs when the effective concentration matches the dissociation constant of the weakest interaction. However, we found that the linker-length scaling of avidity cannot be described by models assuming passive linkers. This suggests compensating mechanisms involving long, disordered protein linkers. Understanding these mechanisms is key to understanding avidity in multivalent protein interactions and its role in disorder-based allostery.

## Materials and methods

### Preparation of DNA constructs

Constructs were based on a previously developed biosensor for effective concentrations,^15^ and created by insertion of synthetic genes (Genscript) into either this construct or between the NdeI and BamHI sites of a pET15b vector. The full sequences of the fusion proteins are given in the supplementary materials. Point mutations were obtained using a QuickChange kit (Agilent). Linker regions with different lengths sub-cloned between construct using the NheI and KpnI sites flanking the linker. All constructs were verified using DNA sequencing.

### Protein expression and purification

All fusion proteins were expressed in BL21(DE3) cells in ZYM-5052 auto-induction medium^20^ supplied with 100μg/mL ampicillin and shaking at 120 RPM. For proteins containing fluorescent protein domains, the temperature was kept at 37°C for the first 3 h and thereafter decreased to 18°C. The cells were harvested after 48-72 h, when the cultures changed color indicating mature fluorescent proteins. The proteins without fluorescent protein domains were expressed overnight at 37°C.

The bacterial pellets containing fusion proteins were dissolved in binding buffer (20 mM NaH_2_PO_4_ pH 7.4, 0.5 M NaCl, 20 mM imidazole), lysed by sonication and pelleted by centrifugation (15min, 14.000g). For proteins not containing folded domains (P_15_ and ACTR peptide), the cells were lysed by heating to 80°C for 20 min.^21^ All proteins were purified using gravity flow columns packed with nickel sepharose. A/P variants, A_15_, P_15_ and ACTR peptide were eluted by stepwise increase of the imidazole concentration to 0.5 M. M/N variants were eluted with a single step of 0.5 M imidazole and subsequently purified using Strep-Tactin XT Superflow (IBA) columns according to the manufacturer’s instructions. A_15_ and P_15_ were released by overnight TEV cleavage at 4°C, which for P_15_ was followed by removal of the tag by reverse IMAC. For all proteins except the ACTR peptide, the final purification step was gel filtration (Superdex 200 increase 30/100) into phosphate buffer saline (PBS). The ACTR peptide was dialyzed into PBS and concentrated. Samples were stored on ice until analysis. Protein concentrations were measured using A_280_.

### Measurement of effective concentrations

The measurement of the effective concentrations were based on previous protocols developed for a single chain biosensor.^15^ A constant concentration (1 μM) of all nine combinations of M/N and A(L17A)/P were titrated with the ACTR peptide through 16 serial two-fold dilutions in PBS. The starting concentration was 2.7mM for WT ACTR and 2.2mM for L17A ACTR. Samples were analyzed in triplicate in black 386-well plates with 1g/L bovine serum albumin (BSA) added to prevent sticking. The FRET measurements were performed in a SpectraMax I3 plate reader using donor excitation at 500nm, and emission detected in 15nm-wide bands centered at 535 and 600 nm. The titration data was analyzed using the non-linear fitting function of MATLAB using the following equation:

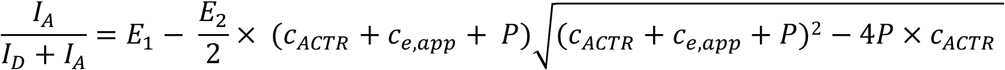

When the titration was done with the WT ACTR peptide this fit determines an “apparent effective concentration” that is corrected by a factor corresponding to the ratio between the apparent effective concentration, and the directly measured effective concentration resulting for titration with the L17A ACTR peptide. The correction factor was established using longest fusion proteins (M/N_120_ and A/P_120_) and applied to all other complexes.

### Surface plasmon resonance (SPR) analysis bivalent complexes

The SPR analysis were performed on Biacore T200 instrument (GE Healthcare) in PBS with 0.1% BSA and 0.05% Tween20. All measurements were performed at 25^°^C with flow rate 30 µl/min. The CMD 500M (Xantec) chip was prepared by amine coupling of anti-mouse IgG antibodies at pH 5.0 (Mouse antibody capture kit, GE Healthcare) according to the protocol described by manufacturer. Immobilization resulted in a capture of approximately 8000 response units (RU) of anti-mouse IgG antibodies on both active and reference flow cell. In each cycle monoclonal mouse anti-Strep-tag II antibody (StrepMAB immo, IBA) was captured on both flow cells to the level of 300 RU. The anti-Strep-tag antibody forms high-affinity complex with SA-Strep-tag (SAWSHPQFEK) and can be used for oriented capture of Strep-tag fused proteins. Between the cycles the chip was fully regenerated with three injections of low pH buffer containing 10 mM glycine pH 1.7.

To determine binding kinetics of M/N:A/P interaction, M/N with C-terminal SA-Strep-tag II was captured on the active flow cell to the level of appr. 100 RU, and A/P or P_15_ was then injected to both flow cells. The concentration of injected protein was varied by a serial two-fold dilution of (A/P:100nM – 0.2nM, P_15_:800nM – 0.7nM). Non-specific binding was removed from the raw binding curves by subtraction of signal from the reference cell of the parallel experiment performed without M/N. Buffer injection was also subtracted from all binding curves. Association was monitored for 60 sec of constant protein injection and dissociation for 120 – 180 sec of constant buffer injection. For the A/P_20_ binding to M/N_20_ combination, the dissociation time was increased up to 15 min as an additional control. Kinetic constants were determined by fitting 1:1 Langmuir interaction model to the binding curves in the Biacore T200 Evaluation Software (GE Healthcare). Analysis of all A/P variants were restricted to concentrations of 25nM and lower. Raw data with the fits is found in Fig. S2-6 and fitted kinetic constants are given in table S2.

### Isothermal titration calorimetry (ITC)

All ITC measurements were performed on MicroCal^TM^ iTC_200_ (GE, Healthcare) at 25°C. For all monovalent interactions, A_15_ variants were titrated into the M/N_20_ using the following concentrations: M/N_20_ 25 µM and A_15_ 250 µM, M/N_20_ 60 µM and A(I34V)_15_ 604 µM, M/N_20_ 40 µM and A(L17A)_15_ 784 µM, M/N_20_ 130 µM and A(L17A/I34V)_15_ 1.31 mM. For bivalent interactions A/P_20_ and A/P_120_ at a concertation 100 µM were titrated into M/N_20_ or M/N_120_ at a concertation 10 µM. Heat of titrant dilution was determined by an analogous titrated into PBS, and was subtracted from the binding data prior to analysis. Data manipulation and analysis were performed in Origin^TM^ (OriginLab Corporation) included in the MicroCal software. For both monovalent and bivalent interactions a single-site model was fitted giving K, ΔH and N value. All measurements, except M/N_20_ with A(L17A/I34V)_15_, were done in triplicates from which mean values with standard deviation were calculated and shown in table S4 and table S5.

### Microscale thermophoresis (MST)

Measurements were performed on Monolith NT.115 (NanoTemper) in PBS using “Standard” capillaries. For determination of A(L17A/I34V)_15_:M/N_20_ binding affinity reaction mixes containing 250 nM of A(L17A/I34V)_15_ and M/N_20_ in 2-fold serial dilution (865 μM – 0.053 nM) were prepared. The reaction mixes were loaded in MST capillaries and analyzed at 20% MST power and a LED intensity of 20% observing the fluorescence from mRuby3. Data analysis was performed in NT analysis 1.5.41 software.

## Results

### Design of model system

To explore the role of linker architecture in avidity, we made two fusion proteins consisting of the two halves of two interaction pairs joined by a disordered linker (Fig. 1B). We used the interaction domains from ACTR:CREB binding protein and p66α:MBD2 such that MBD2 and NCBD are linked to become M/N and ACTR and p66α are linked to become A/P. Together, M/N and A/P form a bivalent complex. We indicate linker length as subscript such that M/N_120_ indicates MBD2 and NCBD linked by a 120-residue linker. The complexes consist of small domains (Fig. 1C)^22,23^, have nanomolar affinities and have been characterized thoroughly by mutagenesis.^24,25^ In the following, we use numbering consistent with the PDB files 1KBH and 2L2L. These interaction domains are bigger than most MoRFs, but allowed us to generate a range of affinities through mutagenesis. To measure the monovalent affinities, we also made the two halves of the proteins, A_15_ and P_15_, each containing 15 flanking disordered residues from the linker. Furthermore, a SA-Strep-tag was included in C-terminus of M/N to allow immobilization for SPR experiments.

To quantify the effect of the linker architecture, we designed the model system to allow direct measurement of the effective concentration of ring closure. Recently, we developed a biosensor that allows measurement of intra-molecular effective concentrations.^15^ We adapted this system by introducing mClover3 N-terminally in A/P and mRuby3^26^ C-terminally in M/N (Fig. 2A). When ACTR and NCBD forms a complex, the FRET pair should come in close contact and allow FRET. Titration of the bivalent complex with ACTR should compete with the intra-molecular complex and force ring opening. To allow the free peptide to compete efficiently, we introduced a mutation in the ACTR domain of A/P (L17A) to weaken the interaction. This shifts the titration midpoint by the affinity difference between L17A and WT ACTR, and this ratio subsequently applied as a correction factor to produce the true effective concentration.

**Figure 2:**
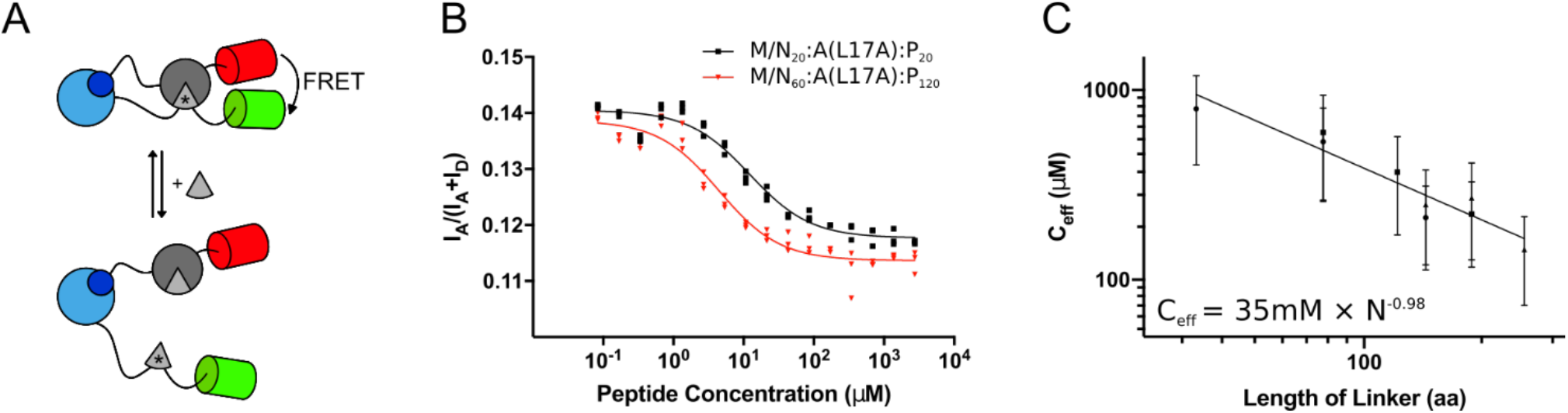
Measurement of effective concentration of ring-closure. (A) The bivalent fusion proteins allow direct measurement of the effective concentrations of ring-closure using competition assays. Free ACTR WT peptide can displace the intramolecular interaction and results in reduced FRET from the nearby fluorescent domains. (B) Titration experiments reveal the apparent effective concentration as the mid-point of the titration curve. As the A(L17A) variant is used in the fusion protein, a correction factor corresponding to the difference in affinity between A(L17A) and WT is applied to produce the true effective concentration. (C) Effective concentrations has a polynomial dependence on total linker length in analogy with scaling laws used in polymer physics.

Linkers can have many different architectures, but perhaps the most general type of linker is a disordered chain of variable length. We recently showed that the sequence of such linkers has a large effect on effective concentrations due to attractive or repulsive interactions.^15^ Linker-linker interactions may play important roles in multivalent disordered proteins, but such interactions are undesirable for the purpose of establishing a theoretical baseline. Ideally, we want linkers that solely act as tunable entropic spacers. Therefore, we used uniform (GS)_x_-repeats of 20, 60 and 120 residues in total as linkers. GS-linkers are highly disordered,^27^ and we recently found that their self-interactions are about as favorable as interactions with the solvent.^15^ While no protein linker is likely to be a true “entropic spacer”, GS-linkers likely represent the closest approximation.

### Measurement of effective concentration

Titration of all combinations of M/N:A/P complexes with WT ACTR peptide resulted in a sigmodal decrease of the proximity ratio (Fig. 2B, S1). This decrease indicated ring-opening due to inter-molecular competition, and suggested that the system worked as intended. By using both the WT and L17A as titrants for one M/N:A/P complex (Fig. 2B, S1), we established the correction factor used to calculate the effective concentration for all other complexes titrated with the WT peptide only. Combination of three linker lengths in each fusion protein resulted in 9 different linker architectures spanning from 40 to 240 disordered residues. The effective concentration decreased monotonically with increasing linker lengths (Fig. 2C, Table S1) spanning a 5.5-fold difference in effective concentrations. In principle, a wider range effective concentration would be desirable, but we ruled out shorter linkers to avoid orientation effects and longer linkers resulted in insoluble proteins.

Geometric considerations suggest that effective concentrations should follow a polymer scaling law with a scaling exponent −3 times that for protein size (*v*).^15^ The error bars in Fig. 2C are substantial, but most of the uncertainty of individual values arise due to propagation of errors and are thus systematic errors that do not affect the scaling exponent. The linear relationship found in Fig. 2C suggested that this was indeed the case for our dataset. However, the scaling exponent of −0.98 was much lower than the numeric value observed when the exact same linker sequence was assayed in an intramolecular biosensor using p66α and MBD2 as the interaction pair (−1.46).^15^ This is not likely to be true for the M/N:A/P complexes as a value below −1 would suggested that it was more compact than a sphere. We defer the discussion of the scaling exponent to the Discussion, and in following simply use the measurements as is.

### Binding kinetics of the bivalent complexes

We chose to measure the avidity effects using surface plasmon resonance (SPR) technique, as low-density surface immobilization of one fusion proteins forces 1:1 binding between the bivalent proteins. This approach thus prevents the competition with higher order complexes, that can plague solution based assays of multivalent proteins. Avidity contains contributions from both the association and dissociation rate-constants. The association rate-constant of a bivalent interaction is the sum of the two monovalent association rate-constants and does not depend on the linker architecture. Furthermore, k_on_ for the ACTR:NCBD is likely to be outside the range that can be accurately measured by SPR. Therefore, we concentrated on k_off_, which is also determined most accurately by SPR. SPR measures a change in refractive index proportional to accumulation of mass on the chip, and can thus not follow ring-closing directly. Dissociation from the cyclical complex requires simultaneous release of both binding sites, and should thus be slower than either monovalent complex. In practical application, avidity is often quantified as affinity enhancement factor (β) defined as the ratio between dissociation constants of bivalent and monovalent interactions.^2^ Since k_off_ contains all the information regarding the linker dependence of avidity, we will define β as the ratio of the dissociation rates of strongest monovalent (p66α:MBD2) and the bivalent complexes. This avoids including the uncertainties from the less well determined k_on_ values into the avidity.

We used SPR to determine binding kinetics of A/P to immobilized M/N (Fig. 3A) for all nine linker combinations. The concentration of the analyte was kept below 25 nM to allow ring-closing to compete efficiently with binding of a second molecule of A/P. Furthermore, the experiment was carried out at high flow rates and relatively low densities of the captured ligand to reduce rebinding artefacts. Under these conditions, all binding reactions could be fitted to a 1:1 interaction model (Fig. 3B, S2). A priori, an additional fast dissociation component could be justified corresponding to A/P that is singly-bound at the beginning of the dissociation phase. However, for each complex the dissociation was fitted well by single exponential dissociation with a global off-rate. This simpler analysis produced essentially identical values as more complex model used by the Biacore analysis software, suggesting that neither mass transport effects nor singly-bound species contributed appreciably to the determined rates (Fig. S6A). Due to the slow dissociation, it was impractical to observe the full dissociation for all complexes. Single-exponential dissociation reaction can be fitted from only 5 % total amplitude, but to ensure the robustness of the data we extended the dissociation time to 15 min for the tightest complex (Fig. S7). The longer dissociation phase did not affect the appearance of the trace or the fitted values appreciably, we thus used the shorter dissociation time for the remaining variants. Analysis of the confidence contours suggest that the dissociation phase is well-determined by the data (Fig. S6B). The non-covalent immobilization strategy (Fig. 3A) entails a risk of M/N dissociation from the antibody contributing to the observed dissociation kinetics. To exclude this possibility, we recorded a long dissociation of M/N alone (Fig. S8). The dissociation from the antibody was ~500-fold slower than the most stable bivalent complex, and thus did not affect the measured k_off_. In total, this confirmed that our measured k_off_ values are robust. The fitted association rate constants are more than an order of magnitude slower than those determined previously for ACTR:NCBD by stopped-flow.^28^ As we are likely at the limit for where SPR can accurately probe association kinetics, we do not interpret these values.

**Figure 3:**
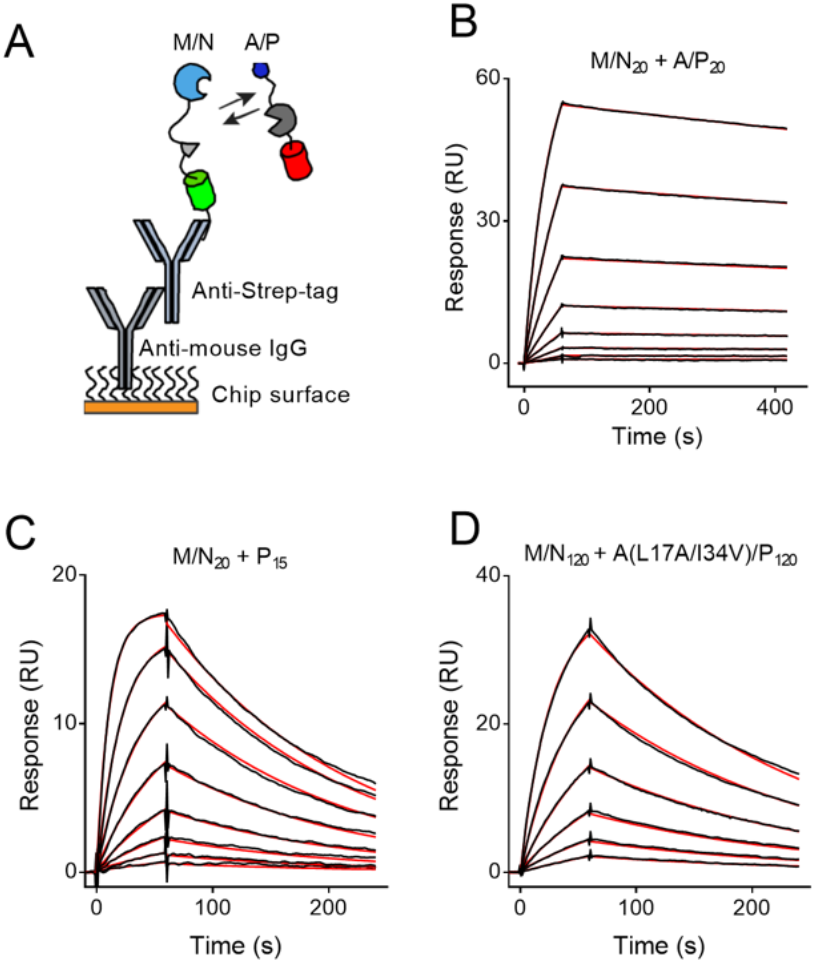
Binding affinity of bivalent interactions measured by SPR. (A) The SPR setup consists of polyclonal anti-mouse IgG antibody directly immobilized on the surface of the SPR chip. Anti-mouse IgG allows oriented capture of monoclonal mouse anti-Strep-tag antibody used for capture of SA-Strep-tag containing M/N variants. Association of A/P to M/N is monitored by injection of single A/P concentration over the chip surface whereas dissociation is followed by constant buffer injection. Example of SPR data for (B) the strongest bivalent (M/N_20_:A/P_20_), (C) monomeric P_15_, and (D) the weakest bivalent interaction (M/N_120_:A(L17A/I34V)/P_120_). Each line represents single A/P concentration from 2-fold serial dilution starting from 25 nM. Raw data is shown in black with the fit shown in red.

The avidity can be quantified by comparison between mono- and bivalent dissociation kinetics. We measured the kinetics of the monovalent variant, P_15_ (Fig. 1B), to all linker variants of M/N. All variants were well fitted by a 1:1 binding model with K_D_ of approximately 3 nM and k_off_ of 6.2 ms^−1^ (Fig. 3C). The independence of the linker length suggest that the linkers do not affect the interaction between the protein binding domains. Comparison of the dissociation phase of bivalent and monovalent complexes (Fig. 3B,C) revealed avidity enhancement in the bivalent system corresponding to a β of 22 and 11, for the shortest and longest linker combinations, respectively. We thus sought to weaken the monovalent interactions to the point where we no longer see avidity enhancement. Based on previous mutagenesis studies,^25^ we introduced mutations into ACTR to reduce the affinity of the weakest interaction. Of the two mutations chosen, one had hardly any effect on the avidity (I34V) whereas the L17A mutation decreased β to about 5 in the complex with the shortest linkers. Combination of these mutations further reduced β to 2.7 and 1.2 for the shortest and longest linkers, respectively. We have thus decreased the avidity by about 10-fold and reached the regime of practically no avidity.

### Affinity of the monovalent interaction

To interpret the avidity enhancement in the bivalent complexes, we needed to determine the NCBD:ACTR affinity. We initially tested the binding of the A_15_ variant to M/N using SPR. These experiments showed binding, but the dissociation phase was too rapid to allow quantification. Instead, the affinity of the monovalent ACTR:NCBD was determined by ITC. All variants of A_15_ could be fitted by a one site model and resulted in binding stoichiometries close to 1 (Table S4). The binding affinity was 0.81 µM for the WT ACTR, 2.58 µM for I34V and 55.7 µM for L17A mutant (Fig. 4A-C, Table S4). The affinity of the interaction between NCBD and ACTR is lower than reported previously,^22^ likely due to small difference in the constructs. We could not achieve precise determination for the double-mutant by ITC due to the large heat of dilution at high concentrations of protein. Instead we resorted to microscale thermophoresis observing the diffusion of the fluorescent protein (Fig. 4D). We could not fully saturate the binding within the limits of solubility, and thereby accurately determine the equilibrium constant. Qualitatively, the data seems roughly consistent with a K_D_ in the hundreds μM, which is in agreement with the predicted K_D_ of 177 µM for the double mutant under the assumption that the mutations are additive.

**Figure 4:**
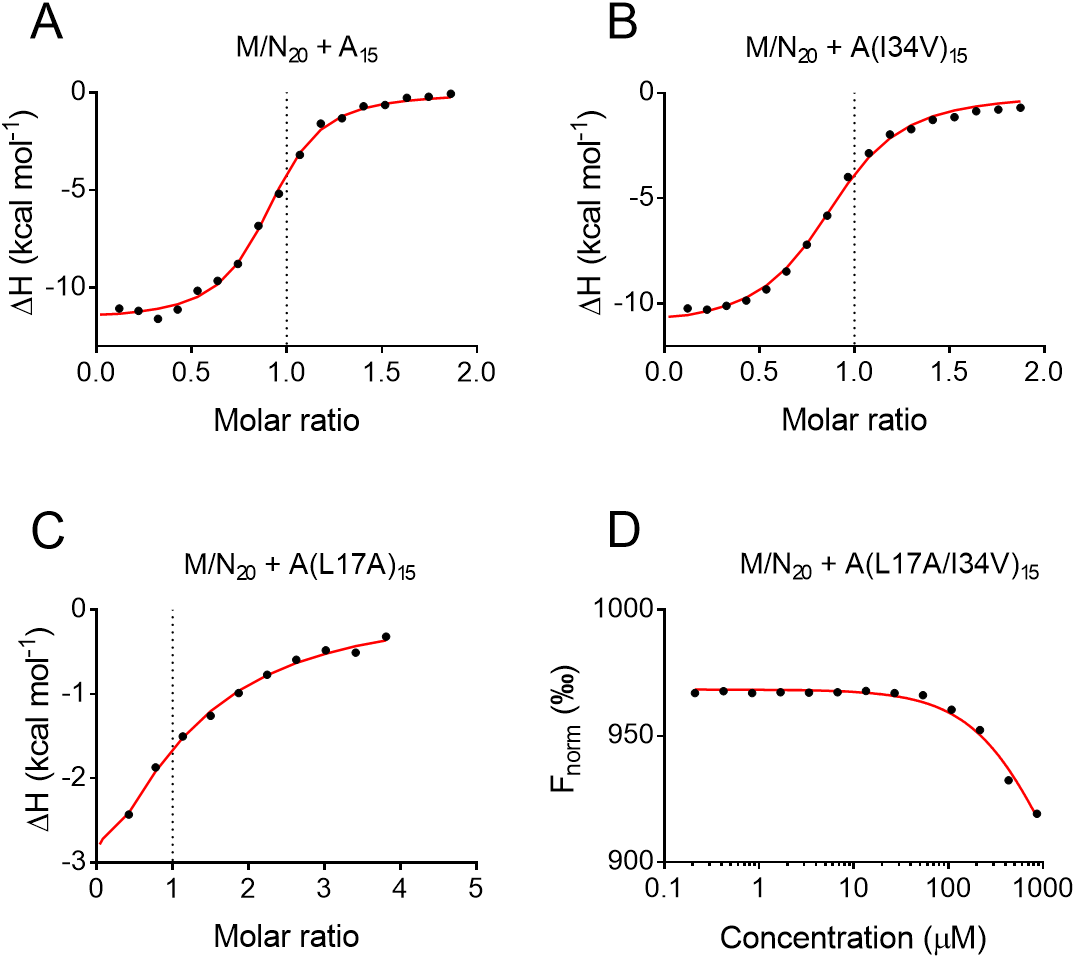
Binding affinity of monovalent interactions. ITC was used to determine K_D_’s for M/N_20_ binding to (A) A_15_, (B) A(I34V)_15_ and (C) A(L17A)_15_. (D) For M/N_20_ and A(L17A/I34V)/P_15_ the K_D_ could not be determined by ITC and was instead estimated by MST. Experimental data is shown in black with the fit shown in red.

### Length scaling of avidity

Avidity enhancement should scale linearly with the effective concentration, which in turn follows polynomial scaling law. In all variants, the avidity enhancement decreased monotonously with linker length as expected (Fig. 5A). The relatively small spread of β-values and the minimal curvature complicated comparison to specific functional forms. We consistently observed an ~2-fold change in the dissociation rates between the shortest and longest linker combinations for all four interaction strengths. The 5.5-fold change in effective concentration between the longest and shortest linkers should thus translate into a 5.5-fold change in dissociation rate and thus β. We thus see a much smaller linker length dependence than expected from effective concentrations. This could be explained if the linkers were not truly entropic spacers, but underwent an attractive enthalpic interaction that partially compensated for the lowered effective concentration. To test this, we compared the binding of the bivalent complexes with 20 and 120-residue-long linkers by ITC (Fig. S9). This interaction was so tight that the only parameter that is robustly determined is the enthalpy. We found that the longest linkers had a less favorable interaction enthalpy than the shortest. This suggest that enthalpic interactions between linkers cannot make up for the discrepancy between the expected and measured scaling with linker length. Early models of avidity suggested that the physical connection was best understood as a coupling energy that is independent of the nature of the binding interaction.^10^ This predicts that a logarithmic plot of dissociation rate-constants should display a constant shift of each variant in good agreement with fig. 5B. Notably, this model assumes full ring-closing and is thus not expected to work near the onset of avidity as is the case for the weakest interaction.

**Figure 5.**
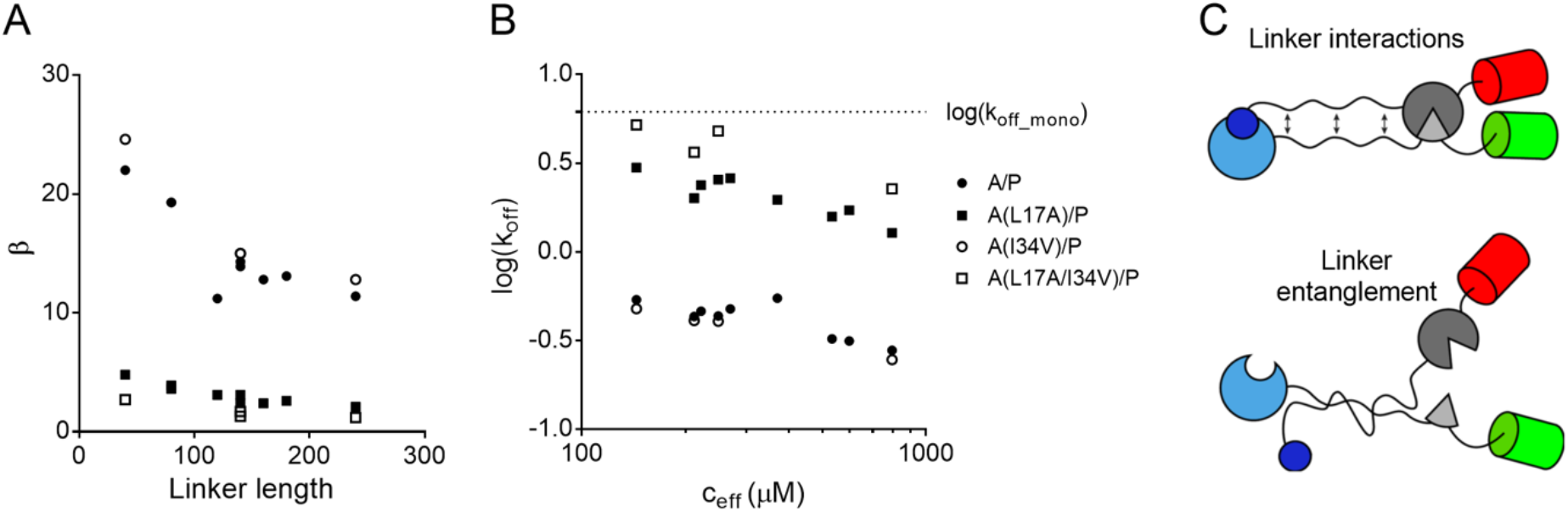
Dependence of avidity on linker length and effective concentration. (A) The avidity enhancement β scales monotonically with linker length consistent with a direct dependence on the effective concentration. (B) The bivalent k_off_ scales identically with c_eff_ for all strengths of the monovalent interaction, consistent with a constant coupling energy for the linkages even near the onset of the avidity. (C) Avidity scales less with linker length than predicted from effective concentrations, which suggests a compensating mechanism that preferentially stabilize fusion proteins with long linkers. We propose two potential mechanisms: Favorable linker-linker interactions, which would have to be entropic to be compatible with the ITC data. Alternatively, the constructs could form complex topologies where the long disordered linkers get entangled and thus remain associated when both binding sites dissociate.

## Discussion

We have generated a model system for studying how avidity depends on the connection between binding sites in disordered and multivalent proteins. The system is artificial, but recapitulates features of many proteins. Studying avidity in an artificial model system has two advantages: First, the connection can be varied in a controlled manner by exchanging the linkers. Second, we can directly measure how the changes in linker architecture affect the effective concentration. Neither of these would be possible in a case-study of a natural protein. Effective concentrations are rarely measured directly in multivalent proteins, but instead estimated from polymer models.^29–33^ The effective concentrations are sensitive to the linker sequence,^15^ so without direct measurement of the effective concentrations it would be difficult to detect deviations from theoretical predictions. We thus believe that the primary strength of the present study is the ability to critically assess theoretical models of avidity in dynamic multidomain proteins.

### Scaling exponent for effective concentrations

The effective concentrations measured here depend less on linker length than expected. When probed in single-chain version of the biosensor, the same linker results in a scaling exponent of −1.46^15^ as opposed to the value of 0.98 found here. Theory suggest that the scaling exponent for effective concentrations should be ~-3*v*, where *v* is the scaling exponent for protein size. The former exponent agrees with theoretical predictions for the chain compaction (*v*=~0.49), and the deviation observed here was thus unexpected and unphysically low. One difference between the two experiments is the presence of the p66α and MBD2 complex inside the linker used here. The effect of a folded domain in the linker, is however expected to be minimal as N- and C-termini are very close in this complex (Fig. 1C). Therefore, this complex only adds a short rigid spacer inside the linker. We do not believe that this is sufficient to cause such a discrepancy. Instead, it is likely due to the different domains used in the competition experiment. The proteins used in the competition experiment reported here (ACTR and NCBD) fold upon binding.^22^ The free ACTR domain is almost entirely unfolded with some propensity to form the first helix.^34–36^ The free NCBD domain is a molten globule with a high degree of complex-like structure.^34,37,38^ The direct comparison of scaling exponent between effective concentration measurements and protein size requires the binding interaction to be approximated by a rigid body docking. This is a bad assumption for proteins that fold upon binding. In such proteins, the disordered parts of the binding domain contribute to the “effective” linker region. As this occurs during the intra-molecular binding reaction, the linker extension effect can likely be by the transition state of the association. The interaction between ACTR:NCBD is initiated by the association between the first partially folded helix of ACTR with NCBD.^25,39^ The N-terminus of NCBD adds about 5 flexible residues to the linker in M/N, whereas the C-terminal segment of ACTR adds about 35 A/P. When the fit in Fig. 1C is repeated with these 40 extra flexible residues added to the linker length, the scaling exponent is increased ~-1.4, in reasonable agreement with results from the single-chain biosensor. Intriguingly, this suggest another mechanism by which coupled folding and binding affects the functional properties of a protein. This will particularly important for proteins with short linkers, where contributions from disordered binding domains may noticeably extend the effective linker. Theoretical predictions of effective concentrations should take structural changes in the complex into account.

### The onset of avidity

Non-covalent interactions form a continuous scale of affinity with no clear lower cutoff. It is thus not clear when a weak interaction ceases to be biologically meaningful. A typical criterion would involve comparison between the K_D_ and the cellular protein concentration as this predicts the whether an interaction is formed. Most MoRFs fail this criterion. Avidity effects can explain how weak MoRFs can be biochemically significant even if their K_D_ is way above the concentration of the protein: An additional weak binding site can act by strengthening another interaction. This suggests an alternative affinity cutoff for MoRFs, where the threshold is based on whether the MoRFs can affect avidity in multivalent complexes. This poses the question of how weak additional interactions elicit avidity. We confirm the theoretical prediction that the onset of avidity occurs when the effective concentration of ring-closing reaches the affinity of the weakest monovalent interaction. The effective concentrations in multidomain proteins typically reach hundreds of µM to low mM.^40^ An avidity-based threshold, thus suggests that a MoRFs should have a K_D_ of 1mM or lower to be considered relevant.

### Deviations from simple models of avidity

Theory suggests that a 5.5-fold increase in effective concentration should result in a 5.5-fold decrease in the K_D_.^16^ In contrast, we only observe a ~2-fold difference in the off-rate between the shortest and longest linker combination. This suggests compensating mechanisms that preclude approximation of even the simplest linker by an entropic spacer. One possible explanation is favorable interactions between the linkers, which would increase with the linker length and compensate for the decrease in effective concentration (Fig. 5C). The scaling exponents observed previously for the GS-linker suggest that our experiments are conducted close to its θ-conditions (*v*=0.5), where intrachain and solvent interactions are equally favorable.^41^ Therefore, the GS-linker is as close an approximation to a passive tether as possible. Nevertheless, ITC suggested that linker expansion is enthalpically unfavorable, which implies that the favorable entropic effect should be even greater than suggested from effective concentrations.

Disordered linkers may regulate interactions in a fundamentally different way, when a system contains more than one such linker. Previously, passive linkers have successfully modelled the role of several protein linkers.^31,42^ Generally, these systems have either been intra-molecular and have only contained one disordered linker. In systems with one disordered linker, the interactions within the linker are adequately described by the scaling exponent of a passive chain. Thus, if you get the dimensions right, you should also get its thermodynamic effects right. In systems with two disordered linkers, interactions between the linkers will add to the stability of the complex. Even measurement of the dimensions of the linker or the effective concentration will not guarantee prediction of the total affinity. The potential for linker interactions will increase with chain length (Fig. 5C), which thus provides the compensating mechanism that reduces the length scaling of avidity. Furthermore, systems with two disordered linkers can experience new types of entropic stabilization such as e.g. chain entanglement (Fig. 5C). A bivalent interaction with long linkers provide plenty of opportunity for chain crossings, which will allow the proteins to stay associated even if both interaction have detached. The complexity of the chain topology will increase with chain length, and thus also provide a compensation in the linker length scaling. Therefore, the role of disordered linkers in the control of biomolecular interactions become significantly more complex when there is more than one, and this likely requires new theoretical description that accounts for the interactions between the linkers.

### Disordered linkers and an expanded concept of allostery

Multidomain proteins are likely to be rich in allosteric regulation. This allows regulatory events such as ligand binding or post-translational modifications to affect interactions through-out the protein. This type of allostery cannot be understood through from single structures as they involve changes in the ensembles of structures occupied.^43,44^ Avidity due to a changed linker architecture is a generic mechanism for regulation by remote effectors, and understanding how linker architecture affects avidity is thus crucial for understanding disordered-based allostery. The reduced length-scaling observed here, suggests that the change in the linker architecture has to be greater than anticipated to translate into functionally important changes in effective concentrations through simple effects on the effective concentration. For long linkers, the most likely source of such regulation would be a change in the charge patterning in the linker, which may have a large effect on linker properties.^45^ On the other hand, since linkers in multi-linker systems cannot be well described by simple passive tether models, this suggest more complex regulatory roles of disordered linkers, where the weak interactions between disordered chains need to be explicitly considered. In total, this expands the scope for how disordered linkers can regulate biochemical properties.

## Acknowledgements

This work was supported by grants to M.K. from the “Young Investigator Program” of the Villum Foundation, Independent Research Fund Denmark (FTP) and the AIAS COFUND program (Agreement No. 609033). We wish to thank Per Jemth and Xavier Warnet for critical comments to this manuscript, and Tanja Klymchuk for technical assistance.

