## Supplemental information for "Linker dependence of avidity in multivalent interactions between disordered proteins"

### Protein sequences of constructs used:

**Color code for features:**
TEV protease site
SA-Strep tag
mClover3
mRuby3
Unique restriction sites
Mutation sites

**Interaction partners:**
MBD2
NCBD
ACTR
p66α

## M/N:
MGSSHHHHHH SSGLVPRGSH MENLYFQSKA FIVTDEDIRK QEERVQQVRK KLEEALMADAS
(Glycine serine linker of variable length)
GTPNRSISPS ALQDLLRTLK SPSSPQQQQQ VLNILKSNPQ LMAAFIKQRT AKYVANQPGM QTSQSQSQSQ
SQSMVSKGEE LIKENMRMKV VMEGSVNGHQ FKCTGEGEGR PYEGVQTMRI KVIEGGPLPF AFDILATSFM
YGSRTFIKYP ADIPDFFKQS FPEGFTWERV TRYEDGGVVT VTQDTSLEDG ELVYNVKVRG VNFPSNGPVM
QKKTKGWEPN TEMMYPADGG LRGYTDIALK VDGGGHLHCN FVTTYRSKKT VGNIKMPGVH AVDHRLERIE
ESDNETYVVQ REVAVAKYSN LGGGMDELYK QSQSQSDYKD DDDKSAWSHP QFEK

## A/P:

MGSSHHHHHH SSGLVPRGSH MVSKGEELFT GVVPILVELD GDVNGHKFSV RGEGEGDATN GKLTLKFICT
TGKLPVPWPT LVTTFGYGVA CFSRYPDHMK QHDFFKSAMP EGYVQERTIS FKDDGTYKTR AEVKFEGDTL
VNRIELKGID FKEDGNILGH KLEYNFNSHY VYITADKQKN CIKANFKIRH NVEDGSVQLA DHYQQNTPIG
DGPVLLPDNH YLSHQSKLSK DPNEKRDHMV LLEFVTAALE SGGEDPMVST GQSQSQSQSQ SEGQSDERAL
LDQLHTLLSN TDATGLEEID RALGIPELVN QGQALEPKAS
(Glycine serine linker of variable length)
GTTSPEERER MIKQLKEELR LEEAKLVLLK KLRQSQIQKE ATAQK

## A15:
MGSSHHHHHH SSGLVPRGSH MVSKGEELFT GVVPILVELD GDVNGHKFSV RGEGEGDATN GKLTLKFICT
TGKLPVPWPT LVTTFGYGVA CFSRYPDHMK QHDFFKSAMP EGYVQERTIS FKDDGTYKTR AEVKFEGDTL
VNRIELKGID FKEDGNILGH KLEYNFNSHY VYITADKQKN CIKANFKIRH NVEDGSVQLA DHYQQNTPIG
DGPVLLPDNH YLSHQSKLSK DPNEKRDHMV LLEFVTAALE SGGEDPMVST GQSQSQSQSQ SEGQSDERAL
LDQLHTLLSN TDATGLEEID RALGIPELVN QGQALEPKAS GSGSGSGSGS GSGSGENLYF QSGSGSGSGS
GSGSGSGTTS PEERERMIKQ LKEELRLEEA KLVLLKKLRQ SQIQKEATAQ K

## P15:
MGSSHHHHHH SSGLVPRGSH MENLYFQSYG SGSGSGSGSG SGSGSGSGSG TTSPEERERM IKQLKEELRL
EEAKLVLLKK LRQSQIQKEA TAQK

#### ACTR peptide:

MGSSHHHHHH SSGLVPRGSH MQSQSQSQSQ SEGQSDERAL LDQLHTLLSN TDATGLEEID RALGIPELVN
QGQALEPKAS GSGSGSGSGS Y

### Supplementary figures:

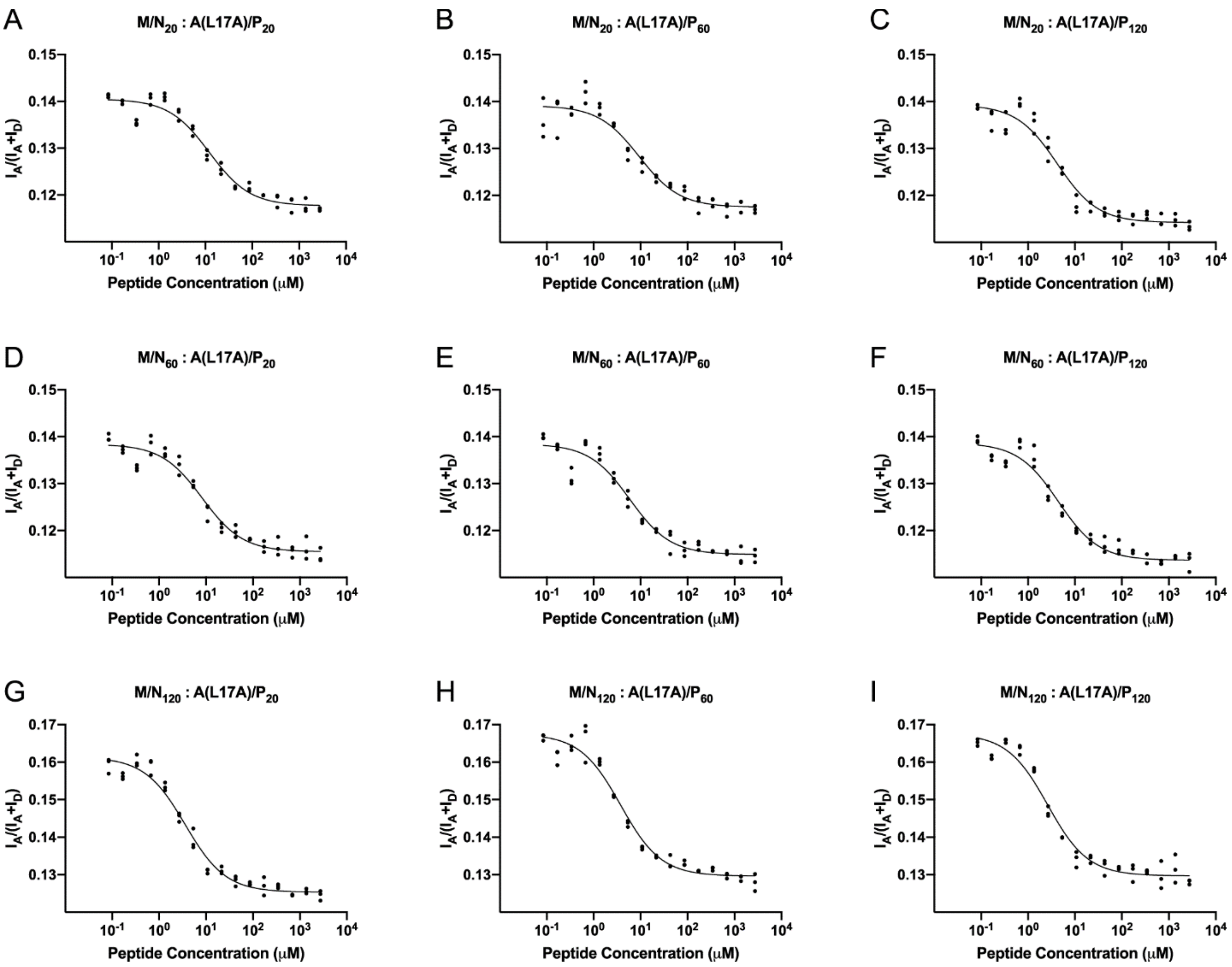


**Figure S1**. Titration data with the fit for the effective concentration measurements. All measurements were performed for M/N and A(L17A)/P variants with various linker lengths. Bivalent interactions were outcompeted by titration of free WT ACTR peptide resulting in a decrease in the FRET signal. Each plot combines raw data from three independent measurements. Combined data was used for the fitting.

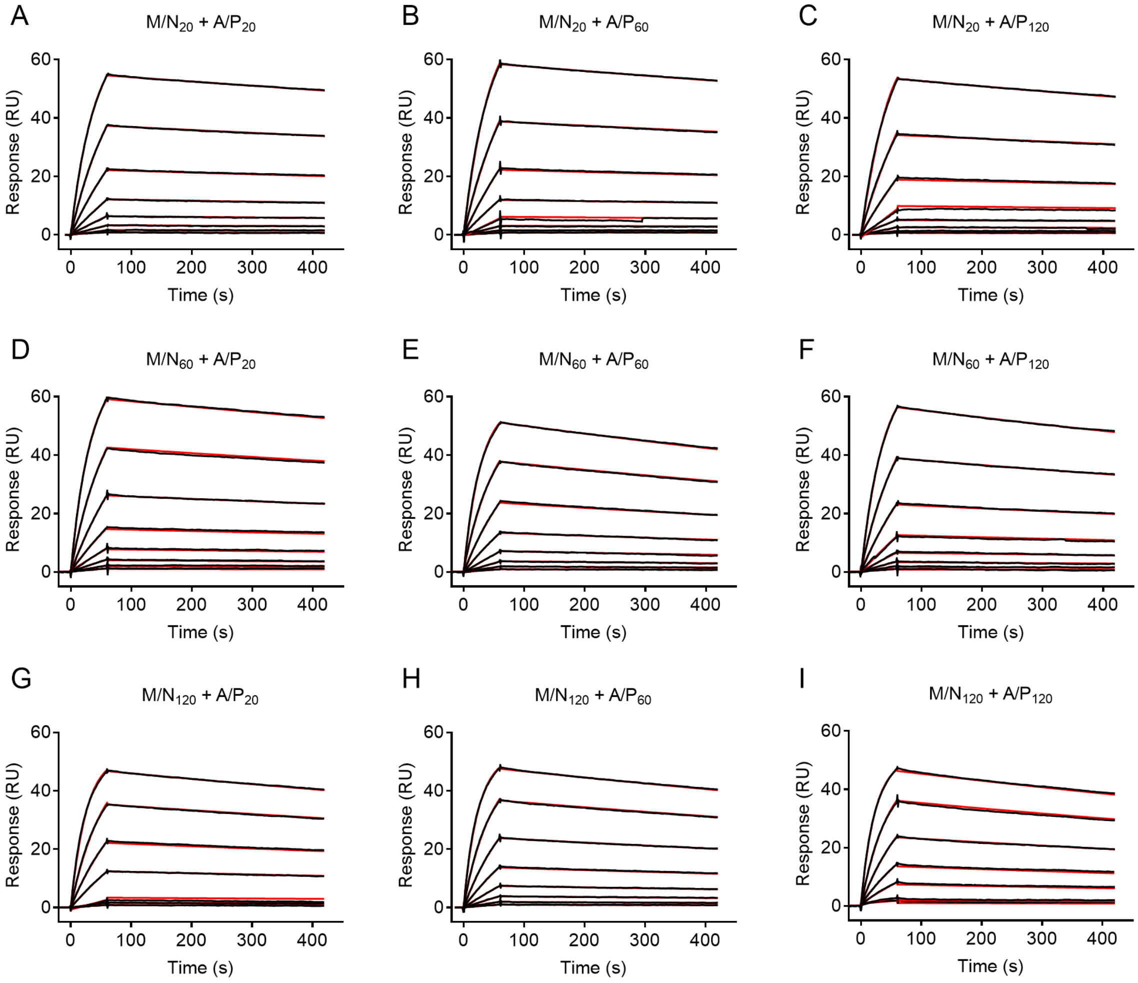


**Figure S2**. Sensorgrams of A/P binding to M/N captured on the chip surface. Each line represents single A/P concentration from 2-fold serial dilution starting from 25nM. Black lines show the raw data, whereas red lines represent the fit of 1:1 binding model.

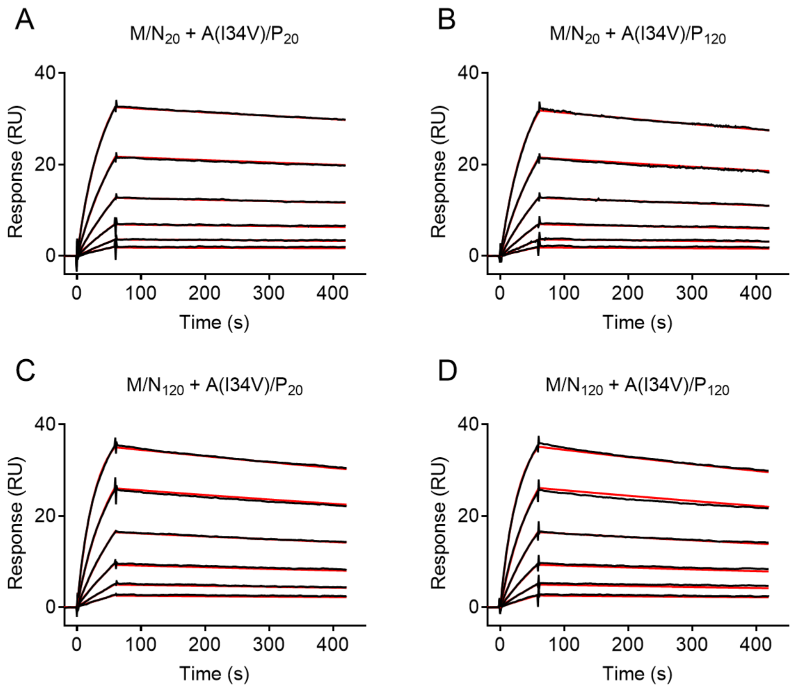


**Figure S3.** Sensorgrams of A(I34V)/P binding to M/N captured on the chip surface. Raw data from 2-fold serial dilution of A(I34V)/P at concentrations 25nM and lower is shown in black with a 1:1 binding model fit shown as red lines.

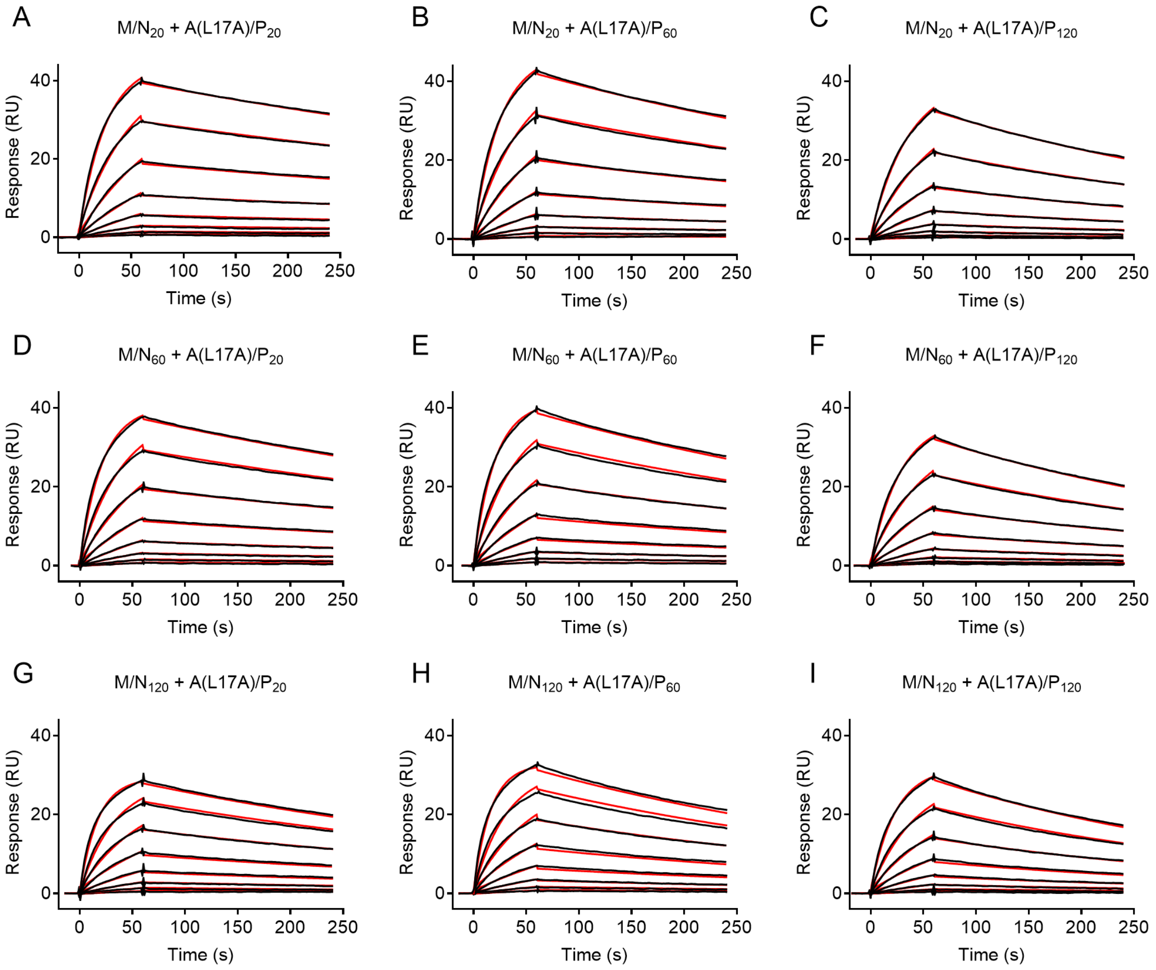


**Figure S4.** Sensorgrams of A(L17A)/P binding to M/N variants captured on the chip surface. Raw data from 2-fold serial dilution of A(L17A)/P at concentrations 25nM and lower is shown in black with a 1:1 binding model fit shown as red lines.

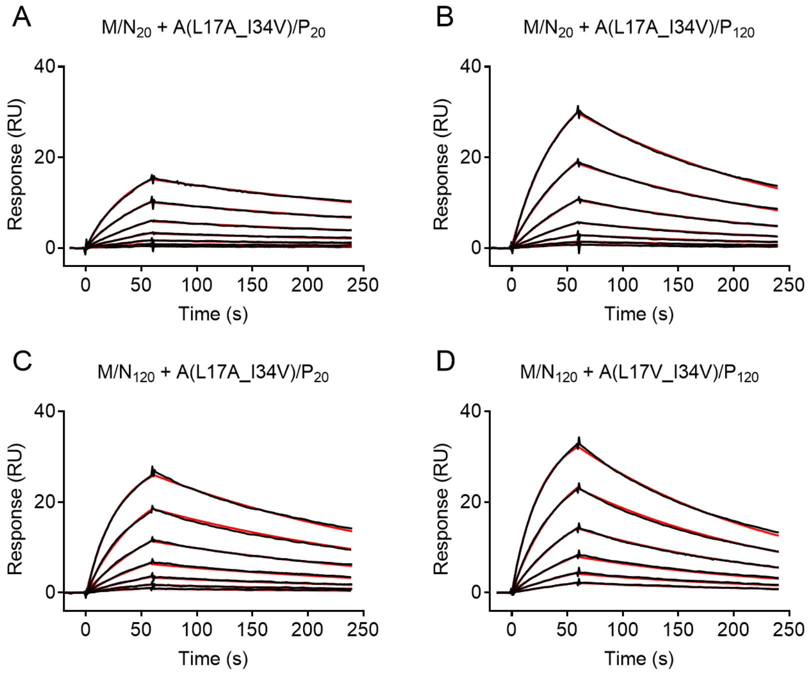


**Figure S5.** Sensorgrams of A(L17A/I34V)/P binding to M/N captured on the chip surface. Raw data from 2-fold serial dilution of A(L17A/I34V)/P at concentrations 25nM and lower is shown in black with a 1:1 binding model fit shown as red lines.

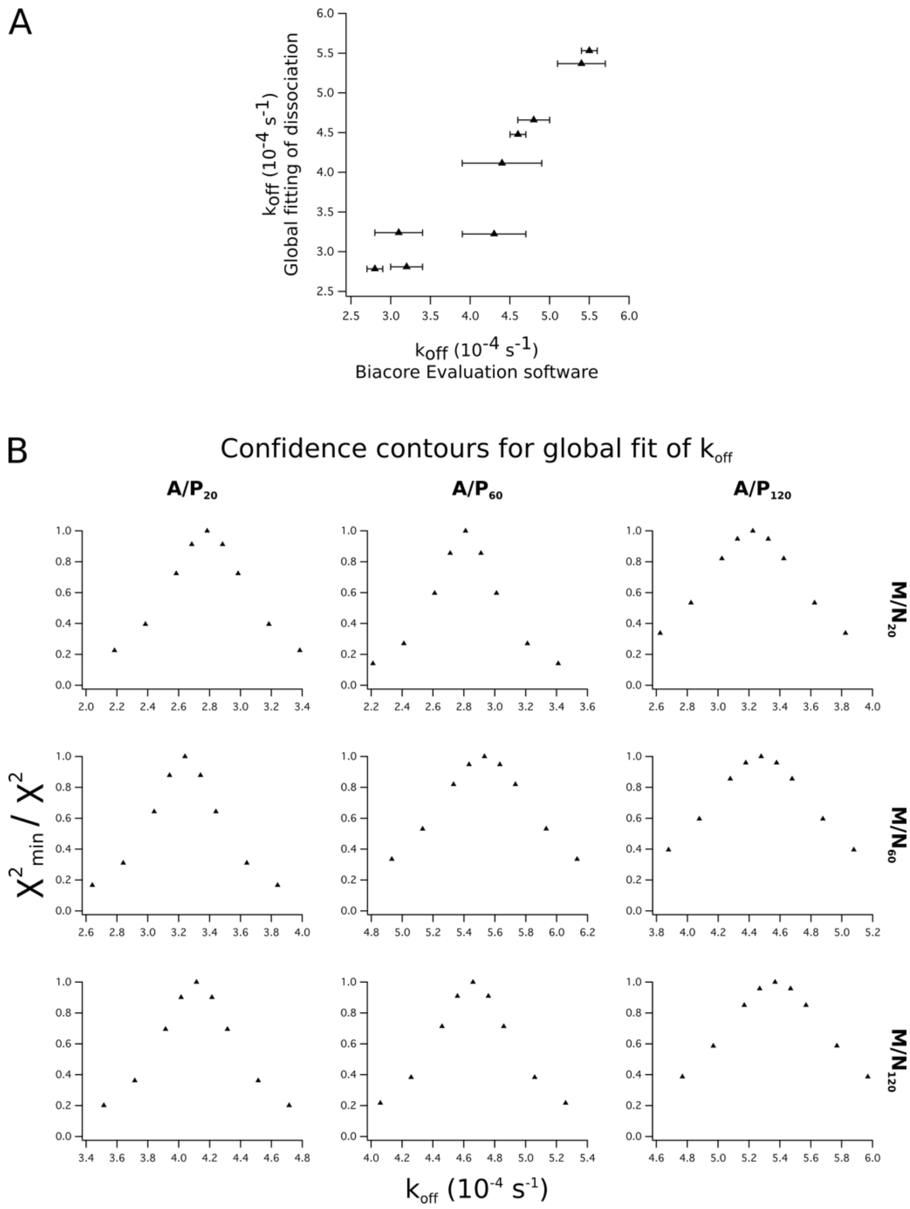


**Figure S6: Confidence analysis of the fitted dissociation rate.** A) The dissociation phase of each complex was analyzed by global fitting to single exponential dissociation reaction by in Igor Pro. This simpler model produced essentially the same dissociation rates as the the Biacore Analysis software. B) Confidence contours for the fitted dissociation rates were analysed in the global fit. For each data set, the dissociation rate occupies a distinct minimum in the $\chi^2$-surface, suggesting that the fitted off-rate is well-constrained by the data.

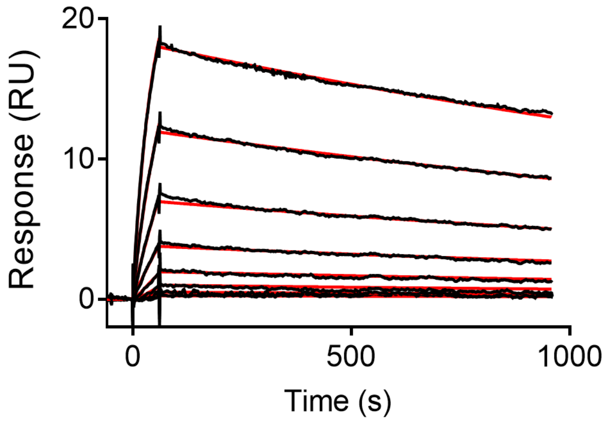


**Figure S7**. Sensorgrams of A/$P_{20}$ binding to M/$N_{20}$ with dissociation time extended to 900 sec. Each line represents single A/P concentration from 2-fold serial dilution starting from 25nM. Black lines stand for raw data, whereas red lines represent the fit of 1:1 binding model. Fitted $k_{off}$ is comparable with data obtained for shorter dissociation time (180 sec) and equals to 0.36 ± 0.01 $ms^{-1}$.

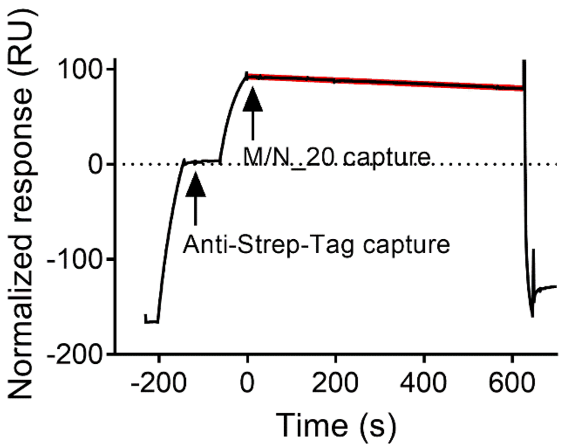


**Figure S8**. Dissociation of M/N from the chip surface. To correct the A/P:M/N dissociation for the loss of M/N due to dissociation from the chip the analogical experiment was done with the constant buffer injection (no A/P). The sensorgram represents signal from accumulation of anti-Strep-tag antibody on the chip surface (-200 sec to -140 sec), followed by 60 sec of stabilization period and 60 sec injection of M/N. Rate of M/N dissociation from the chip was determined by fitting mono exponential decay to the top part of the curve (fit is show in red), giving the $k_{off}$ equal to 0.55 $\mu s^{-1}$.

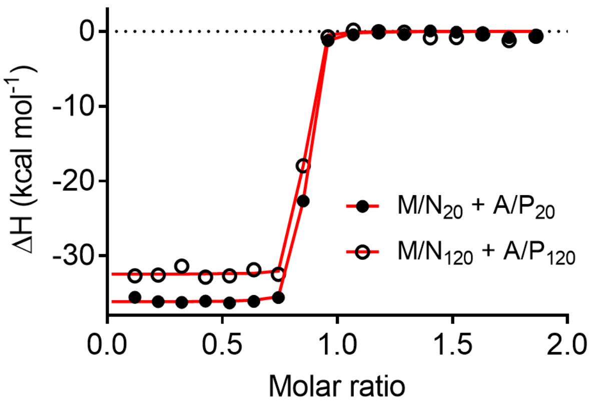


**Figure S9**. Binding of M/N$_{20}$ to A/P$_{20}$ and M/N$_{120}$ and A/P$_{120}$ was followed by ITC. Symbols represent experimental data with the one site model fit shown as a red line.

### Supplementary tables:

**Table S1:**

The effective concentration in μM measured for the different combinations of M/N and A/P.

| Combination | $C_{eff}$ | 95% confidence interval |
|---|---|---|
| M/$N_{20}$ : A(L17A)/$P_{20}$ | 797 | (403-1191) |
| M/$N_{20}$ : A(L17A)/$P_{60}$ | 598 | (259-938) |
| M/$N_{20}$ : A(L17A)/$P_{120}$ | 249 | (120-379) |
| M/$N_{60}$ : A(L17A)/$P_{20}$ | 533 | (263-804) |
| M/$N_{60}$ : A(L17A)/$P_{60}$ | 370 | (172-567) |
| M/$N_{60}$ : A(L17A)/$P_{120}$ | 270 | (127-413) |
| M/$N_{120}$ : A(L17A)/$P_{20}$ | 212 | (112-312) |
| M/$N_{120}$ : A(L17A)/$P_{60}$ | 222 | (116-328) |
| M/$N_{120}$ : A(L17A)/$P_{120}$ | 144 | (73-215) |

**Table S2.**

Kinetic constants determined by SPR for different combinations of M/N and A/P or P_15.

| | $M/N_{20}$ | | | $M/N_{60}$ | | | $M/N_{120}$ | | |
|---|---|---|---|---|---|---|---|---|---|
| | $k_{on}$ $((10^6 {*} M {*} s)^{-1})$ | $k_{off}$ $(ms^{-1})$ | $K_D$ (nM) | $k_{on}$ $((10^6 {*} M {*} s)^{-1})$ | $k_{off}$ $(ms^{-1})$ | $K_D$ (nM) | $k_{on}$ $((10^6 {*} M {*} s)^{-1})$ | $k_{off}$ $(ms^{-1})$ | $K_D$ (nM) |
| $A/P_{20}$ | 1.05 ± 0.006 | 0.28 ± 0.001 | 0.27 | 1.26 ± 0.002 | 0.32 ± 0.002 | 0.26 | 1.78 ± 0.018 | 0.43 ± 0.004 | 0.24 |
| $A/P_{60}$ | 1.32 ± 0.019 | 0.31 ± 0.003 | 0.24 | 1.40 ± 0.001 | 0.55 ± 0.001 | 0.39 | 1.64 ± 0.001 | 0.46 ± 0.001 | 0.28 |
| $A/P_{120}$ | 1.93 ± 0.037 | 0.44 ± 0.005 | 0.23 | 1.28 ± 0.011 | 0.48 ± 0.002 | 0.37 | 1.69 ± 0.003 | 0.54 ± 0.003 | 0.32 |
| $A(I34V)/P_{20}$ | 1.85 ± 0.004 | 0.25 ± 0.003 | 0.13 | n.d. | n.d. | n.d. | 1.97 ± 0.004 | 0.41 ± 0.003 | 0.14 |
| $A(I34V)/P_{120}$ | 2.85 ± 0.004 | 0.41 ± 0.003 | 0.21 | n.d. | n.d. | n.d. | 2.86 ± 0.006 | 0.48 ± 0.004 | 0.17 |
| $A(L17A)/P_{20}$ | 0.75 ± 0.002 | 1.28 ± 0.008 | 1.70 | 0.91 ± 0.002 | 1.58 ± 0.010 | 1.74 | 1.13 ± 0.004 | 2.01 ± 0.016 | 1.78 |
| $A(L17A)/P_{60}$ | 0.76 ± 0.002 | 1.72 ± 0.009 | 3.33 | 0.96 ± 0.003 | 1.97 ± 0.013 | 2.06 | 1.20 ± 0.005 | 2.38 ± 0.019 | 1.98 |
| $A(L17A)/P_{120}$ | 0.51 ± 0.001 | 2.56 ± 0.008 | 4.98 | 0.65 ± 0.002 | 2.61 ± 0.008 | 4.02 | 0.80 ± 0.003 | 2.99 ± 0.014 | 3.72 |
| $A(L17A/I34V)/P_{20}$ | 0.94 ± 0.004 | 2.27 ± 0.011 | 2.41 | n.d. | n.d. | n.d. | 1.27 ± 0.005 | 3.64 ± 0.013 | 2.87 |
| $A(L17A/I34V)/P_{120}$ | 0.83 ± 0.011 | 4.81 ± 0.044 | 5.77 | n.d. | n.d. | n.d. | 1.34 ± 0.005 | 5.20 ± 0.013 | 3.89 |
| $P_{15}$ | 1.68 ± 0.007 | 6.16 ± 0.020 | 3.66 | 2.30 ± 0.016 | 6.01 ± 0.036 | 2.61 | 3.34 ± 0.038 | 6.71 ± 0.067 | 2.01 |

**Table S3.**

Parameters determined by ITC for monovalent interactions between $M/N_{20}$ and $A_{15}$ variants. Values for $A_{15}$, $A(I34V)_{15}$ and $A(L17A)_{15}$ represent mean with standard deviation from three independent repeats, whereas values for $A(L17A/I34V)_{15}$ represent a single experiment.

| | ΔH (kcal/mol) | ΔS (cal/mol/deg) | $K_D$ (μM) | N |
|---|---|---|---|---|
| $A_{15}$ | -13.0 ± 1.1 | -15.6 ± 4.0 | 0.81 ± 0.24 | 0.90 ± 0.03 |
| $A(I34V)_{15}$ | -10.9 ± 0.3 | -10.9 ± 1.2 | 2.58 ± 0.26 | 0.91 ± 0.04 |
| $A(L17A)_{15}$ | -8.3 ± 3.5 | -8.4 ± 12.1 | 55.7 ± 14.8 | 0.87 ± 0.26 |
| $A(L17A/I34V)_{15}$ | -5.04 | 0.56 | 152.9 | 1.59 |

**Table S4.**

Parameters determined by ITC for bivalent interactions between $M/N_{20}$ and $A/P_{20}$ or $M/N_{120}$ and $A/P_{120}$. Values represent mean with standard deviation from three independent repeats.

| | ΔH (kcal/mol) | ΔS (cal/mol/deg) | $K_D$ (nM) | N |
|---|---|---|---|---|
| $M/N_{20}$:$A/P_{20}$ | -36.0 ± 0.2 | -80.2 ± 1.7 | 1.59 ± 1.02 | 0.81 ± 0.002 |
| $M/N_{120}$:$A/P_{120}$ | -33.2 ± 1.4 | -71.3 ± 5.3 | 1.73 ± 0.61 | 0.74 ± 0.06 |
